# DNA methylation in cockroaches is essential in early embryo development and reduces gene expression noise

**DOI:** 10.1101/2020.08.26.267906

**Authors:** Alba Ventos-Alfonso, Guillem Ylla, Jose-Carlos Montañes, Xavier Belles

## Abstract

The influence of DNA methylation on gene behavior, and its consequent phenotypic effects appear to be very important, but the details are not well understood. Insects offer a diversity of DNA methylation modes, making them an excellent lineage for comparative analyses. However, functional studies have tended to focus on quite specialized holometabolan species, such as wasps, bees, beetles, and flies. Here we have studied DNA methylation in a hemimetabolan insect, the cockroach *Blattella germanica*, a model of early-branching insects. In this cockroach, one of the main genes responsible for DNA methylation, *DNA methyltransferase 1* (*DNMT1*), is expressed in early embryogenesis. In our experiments, *DNMT1* interference by RNAi reduces DNA methylation and impairs blastoderm formation. Using Reduced Representation Bisulfite Sequencing (RRBS) and transcriptomic analyses, we observed that hypermethylated genes are associated with metabolism and are highly expressed, whereas hypomethylated genes are related to signaling and have low expression levels. Moreover, the expression change in hypermethylated genes is greter than that in hypomethylated genes, whereas hypermethylated genes have less expression variability than hypomethylated genes. The latter observation has also been reported for humans and in *Arabidopsis* plants. A reduction in expression noise may therefore be one of the few universal effects of DNA methylation.

## Introduction

DNA methylation is the covalent addition of a methyl group to a DNA nucleotide. In most of the animals studied, this only occurs in cytosines, particularly at CpG dinucleotide sites (He et al. 2011; Hunt et al. 2013; Bewick et al. 2017). It is a widespread epigenetic mechanism that contributes to gene expression regulation in eukaryotes (He et al. 2011; Jones 2012; Sarda et al. 2012; Anastasiadi et al. 2018). In mammals, it has been associated with a number of biological processes, including embryo development, genomic imprinting, X-chromosome inactivation, and silencing of retrotransposons (He et al. 2011; Jones 2012).

DNA methylation patterns differ between vertebrates and invertebrates. While in vertebrates DNA methylation is typically localized in the 5’ regulatory regions and appears associated with gene inactivation (Anastasiadi et al. 2018), in invertebrates it is mainly localized in intragenic regions and appears associated with gene activation (Bonasio et al. 2012; Falckenhayn et al. 2013; Hunt et al. 2013; Wang et al. 2013; Glastad et al. 2014; Glastad et al. 2016; Bewick et al. 2019). Moreover, DNA methylation levels in arthropods, particularly insects, are generally lower than in vertebrates (Bewick et al. 2017). Indeed, given the possibilities of comparing different orders that have distinct DNA methylation patterns (Bewick et al. 2017; Lewis et al. 2020a), insects have become the model of choice for studying the functional significance of this DNA modification (Hunt et al. 2013).

Most insects develop through metamorphosis, which can be classified into two modes: hemimetabolan, or direct development through the embryo, nymph, and adult stages; and holometabolan, or indirect development through the embryo, larva, pupa and adult stages (Belles 2020). In this respect, although DNA methylation has been detected in the different insect groups, higher levels have been observed in hemimetabolan than holometabolan models (Falckenhayn et al. 2013; Bewick et al. 2017). This, along with a comparative analysis between hemimetabolan and holometabolan insects, led Ylla et al. (2018) to hypothesize that DNA methylation could be instrumental in the type of embryo development, and the mode of metamorphosis. Many roles have been associated with DNA methylation in insects, including phenotypic plasticity and caste determination (Bonasio et al. 2012; Glastad et al. 2016; Cardoso-Júnior et al. 2017; Li et al. 2018), alternative splicing (Bonasio et al. 2012; Glastad et al. 2014; Glastad et al. 2016), and reproduction (Schulz et al. 2018; Bewick et al. 2019). However, there are few functional studies on the role of DNA methylation during early embryo development (Schulz et al. 2018; Bewick et al. 2019), despite this being the period when de novo DNA methylation is expected to occur (He et al. 2011).

DNA methylation is catalyzed by DNA-methyltransferases (DNMTs). In mammals, DNMTs are classified into DNMT3, which establishes new methylation (methylation de novo), and DNMT1 which preferentially methylates hemimethylated DNA, maintaining methylation during successive cell generations (maintenance methylation) (He et al. 2011; Jones 2012). Although a third DNMT was initially reported, further studies demonstrated that the so-called DNMT2 actually methylates tRNA, rather than DNA (Goll et al. 2006; Jurkowski et al. 2008; Lyko 2018). Insects can possess either just DNMT1 (like the lepidopteran *Bombyx mori* and the coleopteran *Tribolium castaneum*), both DNMT1 and DNMT3 (like the hymenopteran *Apis mellifera*), or neither (like the dipteran, *Drosophila melanogaster*) due to secondary loss of both DNMT1 and DNMT3 (Bewick et al. 2017).

In a previous work, we reported the gene expression patterns in the German cockroach, *Blattella germanica*, on the basis of 11 transcriptomes representing key stages of embryonic and post-embryonic development (Ylla et al. 2018). One of the genes with the most characteristic profile was *DNMT1*, whose expression was concentrated in the first days of embryogenesis. At that time, we hypothesized that DNMT1 would catalyze DNA methylation and that it may play an important role in early embryo development. This work was planned to test these hypotheses, and has enabled us to characterize the DNMT1 gene of *B. germanica.* We have used quantitative PCR to confirm that its expression does indeed concentrate in the first days of embryo development, and that it promotes the methylation of DNA cytosines.

Moreover, maternal RNAi studies have revealed a crucial role of DNMT1 in early embryogenesis. Beyond these findings, by analyzing the genome-wide methylation profiles in regions with a high CpG content, and looking at the relationships between methylation levels and gene expression, we discovered certain regularities that may be of more general interest. Comparing hypermethylated and hypomethylated genes, we found that the former are related to metabolism and are highly expressed throughout development, while the latter are more associated with signaling pathways and generally have low expression levels. Moreover, hypermethylated genes present a relatively high degree of expression change after the DNMT1 peak, but with little expression variability, whereas hypomethylated genes display the opposite properties.

## Results

### *Blattella germanica* has *DNMT1* and *DNMT3* genes that express in the early embryo

By combining a BLAST search in *B. germanica* transcriptomes (Ylla et al. 2018), mapping of the resulting sequences in the *B. germanica* genome (Harrison et al. 2018), and PCR strategies, we obtained a cDNA of 4,662 nucleotides comprising the complete ORF (GenBank accession number MT881788), whose conceptual translation gave a 1554 amino acid sequence that was highly similar to insect DNMT1 proteins. We also obtained a cDNA of 1,803 nucleotides, comprising the complete ORF (GenBank accession number MT881790), whose conceptual translation gave a 601 amino acid sequence that was highly similar to insect DNMT3 proteins. A phylogenetic analysis using DNMT1 and DNMT3 sequences from representative species showed that the DNMT1 and DNMT3 identified in *B. germanica* clustered at the DNMT1 and DNMT3 nodes, respectively (supplementary fig. S1, Supplementary Material online), strongly suggesting that these were DNMT1 and DNMT3 orthologs.

In *B. germanica*, both the *DNMT1* and *DNMT3* genes have a series of introns that could allow the generation of protein isoforms through alternative splicing. However, the analysis of the RNA-seq data (Ylla et al. 2018), together with the fact that we did not amplify more than one product using PCR, robustly supports the idea that there are single DNMT1 and DNMT3 proteins in the *B. germanica* embryo. With regard to protein organization, *B. germanica* DNMT1 contains all the characteristic DNMT1 domains described by Lyko (2018) (fig. 1A): a DNMT1-associated protein 1 (DMAP1) binding domain, which allows interaction with the transcriptional repressor DNMAP1 and the histone diacetylase HDAC2 (HD2); a replication foci targeting sequence (RFTS), which allows DNMT1 to target replication foci; a CXXC domain, which allows DNMT1 to bind unmethylated DNA; two bromo-adjacent homology (BAH) domains, whose function is still unknown; and a catalytic domain at the C-terminal. *B. germanica* DNMT3 also contains all the characteristic DNMT3 domains described by Lyko (2018) (fig.1A): a PWWP domain, which allows binding to histone H3 molecules that are trimethylated at lysine 36; an ATRX-DNMT3-DNMT3L (ADD) domain, which mediates targeting to histone H3 molecules with unmethylated lysine 4; and a catalytic domain, the C5-Cytosine specific DNA methylase domain. In previous analyses, we also found a *bona fide* DNMT2 ortholog (Ylla et al. 2018). However, as DNMT2 is considered a tRNA transferase (Goll et al. 2006), the *B. germanica* ortholog is beyond the scope of this work, which focuses on DNA methylation.

**FIG. 1.**
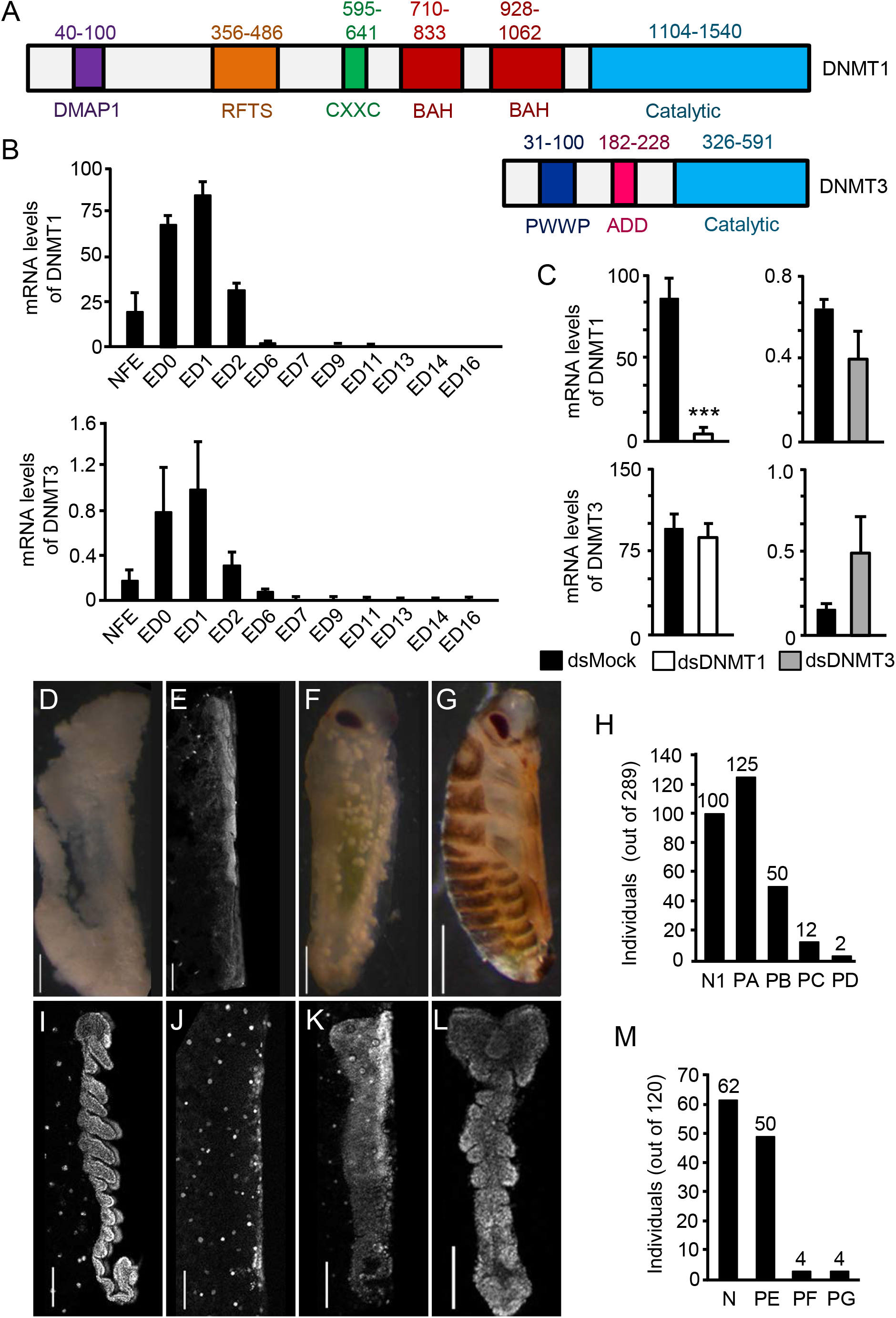
*Blattella germanica* DNMT1 and DNMT3, and effects of maternal RNAi. (A) Protein organization of DNMT1 and DNMT3; the domains follow the nomenclature established by Lyko (2018); numbers indicate the start and end amino acids of the different domains. (B) qRT-PCR mRNA levels of *DNMT1* and *DNMT3* during embryogenesis; NFE: non-fertilized egg; ED0 to ED16: embryo day 0 to embryo day 16; (C) Effects of DNMT1 and DNMT3 maternal RNAi on *DNMT1* and *DNMT3* transcript levels; dsDNMT1, dsDNMT3 or dsMock were injected into 5-day-old adult females, and measurements were taken on ED1. (D-G) Phenotypes observed in unhatched oothecae from DNMT1 depleted embryos; D: Phenotype PA, embryos with development interrupted at the pre-blastoderm stage; E: Phenotype PB, embryos with malformed head and appendage-like structures; F: Phenotype PC, embryos around Tanaka stage 13, with no appendages and a narrower abdomen than normal; G: Phenotype PD, embryos at Tanaka stage 18, ready to hatch, but with darker coloration than normal. (H) Number of individuals showing the phenotypes PA to PD; the total number of individuals studied was 289, and the number of individuals in each category is indicated at the top of each bar; the sample also includes the 100 nymphs hatched from 3 viable oothecae (N1). (I-L) Phenotypes observed in ED4 from DNMT1 depleted embryos; I: Normal ED4 embryo (N); J: Phenotype PE, embryos with development interrupted at Tanaka stage 2; K: Phenotype PF, embryos with development interrupted at Tanaka stage 3; L: Phenotype PG, embryo with a general morphology similar to Tanaka stage 4, with the cephalic and thoracic segments delimited but incompletely developed, and the abdominal region amorphous and unsegmented. (M) Number of embryos showing the phenotypes PN and PE to PG; the total number of embryos studied was 120, and the number of embryos in each category is indicated at the top of each bar. In D-G, and I-L, the upper part of each picture corresponds to the cephalic part of the embryos; the scale bars are equivalent to 500 μm in panels D-G, and 100 μm in panels I-L. In Figures B and C, each qRT-PCR value represents three biological replicates and is expressed as copies of mRNA per 1000 copies of Actin-5c mRNA (mean±SEM); the triple asterisk indicates statistically significant differences with respect to controls (p < 0.001), calculated on the basis of a Pairwise Fixed Reallocation Randomization Test implemented in REST (Pfaffl, 2002).

By using real-time quantitative reverse transcription PCR (qRT-PCR), we studied the expression of DNMT1 and DNMT3 during *B. germanica* embryogenesis (embryo days 0, 1, 2, 4, 6, 7, 9, 11, 13, 14, and 16). The results show that both *DNMT1* and *DNMT3* are expressed between days 0 and 2 of embryogenesis (0 to 12% of embryo development), both showing an expression peak at day 1 (fig. 1B). However, the expression levels of *DNMT3* are very low (maximum expression of 0.98 ± 0.46 copies per 1000 actin copies at day 1), not only when compared with that of *DNMT1* (maximum expression of 86.06 ± 8.83 copies per 1000 actin copies at day 1), but also taking into account the very low quantity of absolute mRNA at this early embryo stage, including the absolute levels of actin mRNA.

### Maternal RNAi of *DNMT1* and *DNMT3*

To study the possible functions of DNMT1 and DNMT3 in the early embryo, we used maternal RNAi. Five-day-old adult females (AdD5) of *B. germanica* were injected with 3 μg of a dsRNA targeting DNMT1 (dsDNMT1) or DNMT3 (dsDNMT3). These females were then allowed to mate (fertilization was checked at the end of the experiment by examining the presence of spermatozoids in the spermatheca), and to produce the first ootheca. Control females were treated equivalently, but with a non-specific dsRNA (dsMock). To estimate the efficiency of the RNAi, we measured the levels in the respective transcripts in 1-day-old embryos, the day of peak expression. In the dsDNMT1-treated females, the mRNA levels of DNMT1 were 87.5% lower than in the controls (fig. 1C), indicating that the maternal RNAi was remarkably efficient. Additionally, the DNMT3 mRNA levels were similar in both groups, indicating that dsDNMT1 is specific and does not affect DNMT3 transcripts. In contrast, dsDNMT3 treatment did not significantly affect the mRNA levels of DNMT3, despite the high dose of dsRNA and the replication using three experimental batches containing 10 females each. Figure 1C illustrates representative results demonstrating that our dsDNMT3 treatments did not reduce *DNMT3* mRNA levels. As we were unable to knock down *DNMT3*, we continued the functional studies with DNMT1.

### Depletion of DNMT1 impairs embryo development

A total of 10 control (dsMock-treated) females formed the first ootheca on day 8 of the adult stage, which hatched 19 days later, giving a total of 373 first instar nymphs (35-40 nymphs per ootheca, on average). The dsDNMT1-treated females (n=10) also produced the first ootheca on day 8, but only 3 out of 10 oothecae (30%) hatched 19 days later, giving a total of 100 first instar nymphs (30-35 nymphs per ootheca). No nymphs hatched from the remaining 7 oothecae (70%) produced by the dsDNMT1-treated females. The examination of the embryos in the 7 unviable oothecae, 20 days after the formation of the ootheca (n=189 embryos) showed various phenotypes, which were classified into the following four categories. Phenotype PA (fig. 1D): embryos with development interrupted in a pre-blastoderm stage, thus, only white yolk was observed. Phenotype PB (fig. 1E): embryos that were completely transparent under the stereomicroscope, but for which 4’, 6-di-amidino-2-phenylindole (DAPI) staining revealed malformations of the head and appendages. Phenotype PC (fig. 1F): embryos at Tanaka stage 13 (Tanaka 1976) (58% embryo development), but with no appendages, and narrower abdomens than the controls. Phenotype PD (fig. 1G): embryos at Tanaka stage 18, thus, just prior to hatching, but presenting a darker coloration than the controls. A total of 125 embryos of the 189 studied showed phenotype PA (66% of the abnormal embryos and 43% of all DNMT1-depleted embryos). Phenotype PB was represented by 50 embryos (26% of the abnormal embryos and 17% of all DNMT1-depleted embryos). Phenotypes PC and PD were the least frequent; 12 embryos presented phenotype PC (6% of the abnormal embryos and 4% of all DNMT1-depleted embryos), and only 2 embryos presented Phenotype PD (1% of the abnormal embryos and 0.7% of all DNMT1-depleted embryos) (fig. 1H).

Since most of the embryos from DNMT1-depleted females died early in their development, coinciding with the temporal expression of *DNMT1*, we repeated the maternal RNAi experiment, but this time we examined the embryos 4 days after oviposition (ED4). We studied 120 embryos from 5 oothecae produced by control (dsMock-treated) females and 120 embryos from 5 oothecae produced by dsDNMT1-treated females. All the embryos from control females (100%) presented the normal aspect of an ED4 embryo, in other words, 20-25% embryo development and Tanaka stage 5–6 (Tanaka 1976) (fig. 1I). A total of 62 out of 120 embryos (51.7%) examined in oothecae from dsDNMT1-treated females, were normal embryos, similar to the controls. The remaining 58 embryos (48.3%) showed several different phenotypes that were classified into three categories, as follows. Phenotype PE (fig. 1J): embryos with development interrupted at Tanaka stage 2, when the germ band is delimited and slightly expanded on both sides (12% embryogenesis). Phenotype PF (fig. 1K): embryos with development interrupted at Tanaka stage 3 (16% development), when the germ band has started to expand on both sides. Phenotype PG (fig. 1L): embryos with a general morphology similar to Tanaka stage 4, at the start of abdominal segmentation and tail folding (17% embryogenesis), but presenting various malformations: cephalic and thoracic segments delimited but incompletely developed; and amorphous and unsegmented abdominal regions. Phenotype PE was represented by 50 embryos (86% of the abnormal embryos and 42% of all the embryos), while phenotypes PF and PG had 4 embryos each (7% of the abnormal embryos and 3% of all embryos, in both cases) (fig. 1M).

### Depletion of DNMT1 reduces CG methylation levels

To assess whether DNMT1 methylates DNA in *B. germanica*, we studied DNA methylation in DNMT1-depleted and control embryos, following the Reduced Representation Bisulfite Sequencing (RRBS) method. For this purpose, we performed RRBS in two different conditions: 4-day-old control embryos (ED4C) and 4-day-old DNMT1-depleted embryos (ED4T), using four biological replicates per condition. We analyzed the levels of methylated cytosines within CG dinucleotides (mCG) in these two conditions in different genomic features. Firstly, we considered all the regions available from the RRBS, then the sequences corresponding to intergenic regions, the genes (the region that is transcribed, including the UTRs), the promoter region (considering an arbitrary length of 2 Kb upstream of the transcription start site), the 5’ UTR, and the 3’ UTR. Moreover, we examined the levels of methylated cytosines in exonic and intronic regions. We considered all exons as a whole (including the 3’ UTR), the first exon, the last exon, all exons except the first and the last, and the exon of monoexonic genes. We performed an equivalent analysis for introns.

Considering the whole gene and different gene features, we observed that the 3’ UTR regions have the higher average levels of methylation (Table 1). Moreover, CG methylation levels are higher in genic regions than in intergenic regions; and within genes these are higher in exonic regions, particularly 3’ UTR regions. In intronic regions, there is a tendency to show higher levels of CG methylation towards the 3’ region (Table 1). Characteristically, the methylation density (fig. 2A) of the different genetic features shows two clear peaks, indicating that they are either very methylated (80-100% methylation), or have very low levels of methylation (0-4%), practically without any intermediate values. RRBS sequencing also revealed that the levels of CG methylation are lower in DNTM1-depleted embryos than in controls, irrespective of the genomic feature examined. In most cases, the reduction was between 50% and 60%. Greater reductions were observed in the first exon (63.8% reduction) and the single exon of monoexonic genes (73.7% reduction) (Table 1). Consequently, DNMT1-depletion modified the bimodal distribution of CG methylation, since a significant proportion of intermediate values appeared, and the peak of high values (80-100% methylation) was reduced (fig. 2A).

**FIG. 2.**
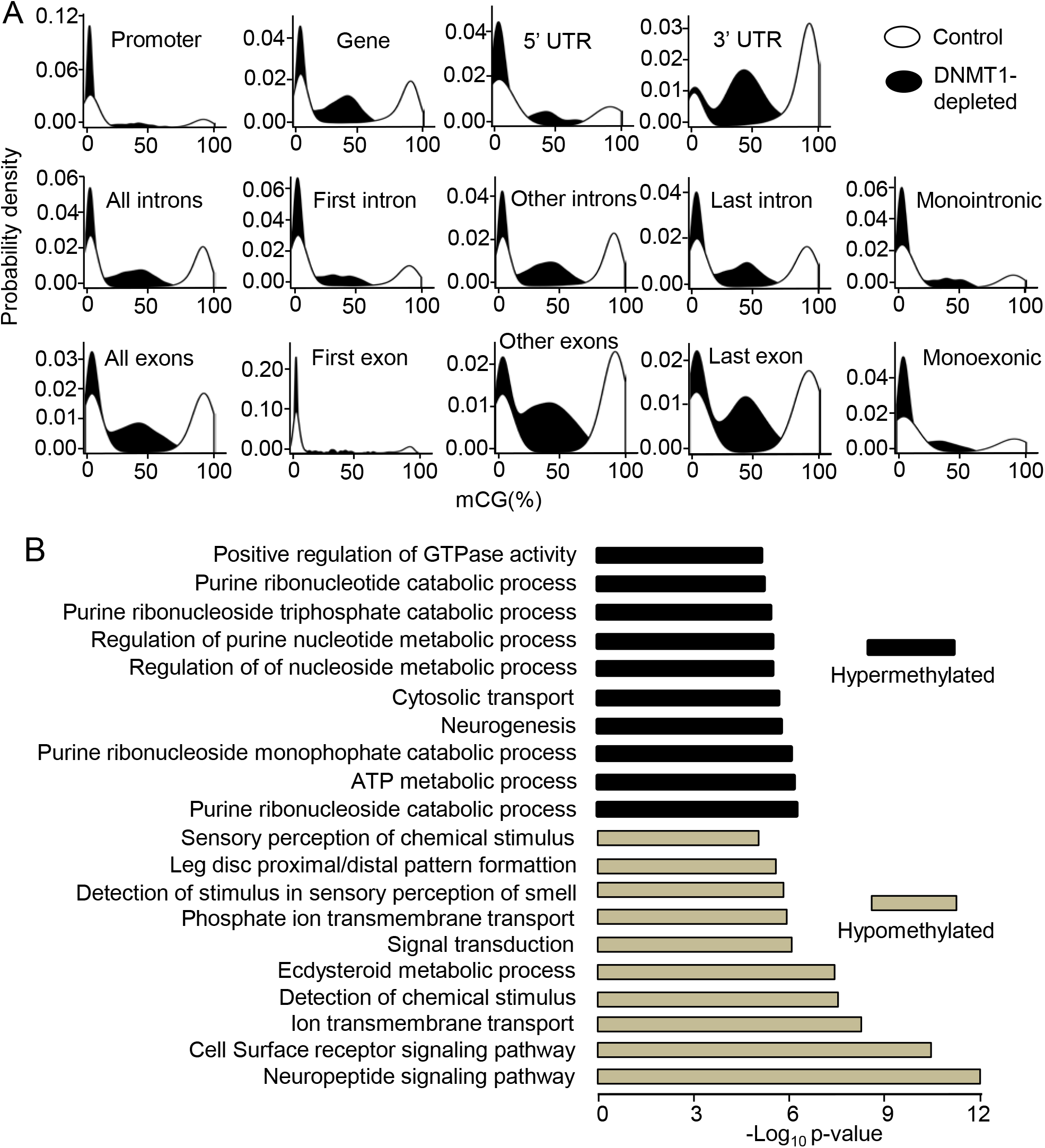
DNA methylation in *Blattella germanica* and effects of DNMT1 depletion. (A) Kernel density plot of CG methylation in control and DNMT1-depleted 4-day-old embryos. The genomic features examined are the same as described in Table 1. In controls the levels of CG methylation are generally either very high (80-100% methylation), or very low (0-4%), thus presenting a bimodal distribution; in DNMT1-depleted insects the bimodal distribution is modified as the peak of high values is reduced. (B) Selection of GO terms of biological functions resulting from enrichment analyses carried out on hypermethylated genes in 4-day-old control embryos. The 10 enriched biological functions with the lowest p-values are shown for hypermethylated and hypomethylated genes. The p-values were calculated according to Fisher’s exact test.

Hypermethylated genes are associated with metabolism and are highly expressed throughout development, whereas hypomethylated genes are associated with signaling and show low expression levels

To obtain information on the functions of the highly methylated genes (80-100% methylation, hereinafter referred to as hypermethylated) and practically unmethylated genes (0-4% methylation, hypomethylated), we carried out a gene ontology (GO) enrichment analysis. The results (fig. 2B) indicate that hypermethylated genes are enriched in biological functions related to metabolic processes, neurogenesis, and cytosolic transport. On the other hand, the potential biological functions of hypomethylated genes appear to be related to signaling pathways, including neuropeptide signaling, cell surface receptor signaling, ion transport, signal transduction, detection of chemical stimulus, ecdysteroid metabolic processes and leg patterning (fig. 2B).

Next, we compared the expression levels of the two types of genes using transcriptomic data corresponding to different stages of *B. germanica* (Ylla et al. 2018). The results show that the expression levels of the hypermethylated genes are higher than those of the hypomethylated genes in all the ontogenetic stages studied, the difference being more evident in early embryo stages, from ED0 to ED6 (fig. 3A).

**FIG. 3.**
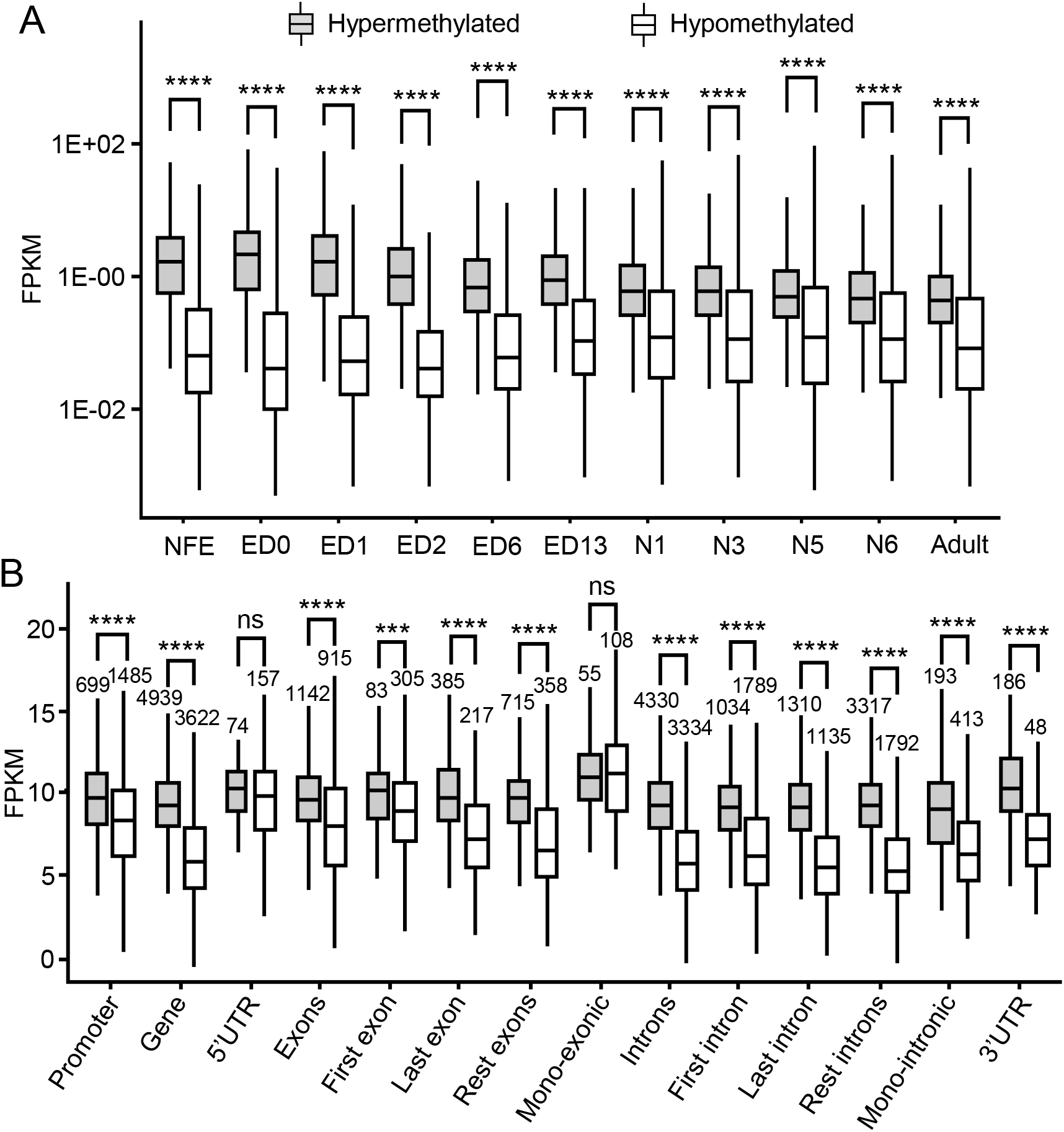
DNA methylation and the amount of gene expression in *Blattella germanica*. (A) Expression levels of hypermethylated and hypomethylated genes in the 11 embryo stages studied (NFE: non-fertilized egg, and ED0 to ED16: embryo day 0 to embryo day 16), four nymphal stages (N1, N3, N5, and N6), and the adult. (B) Expression levels of hypermethylated and hypomethylated genes in 6-day-old embryos (ED6), considering gene expression levels in 6-day-old embryos (ED6), grouped by methylations status (hypomethylated vs hypermethylated) of various gene features. In all cases, expression is expressed as FPKM; the asterisks indicate statistically significant differences using the Mann-Whitney U test, adjusting p-values by False Discovery Rate using the Benjamini-Hochberg method (* FDR < 0.05; ** FDR < 0.01; *** FDR < 0.001; **** FDR < 0.0001), non-significant differences (ns; FDR > 0.05), are also indicated.

We then analyzed the differences in expression between hypermethylated and hypomethylated genes, considering the gene region where the methylation is located. For this analysis, we used the transcriptomic data corresponding to ED6, since this is the stage following the pulse of *DNMT1* expression (fig. 1B). ED6 corresponds to Tanaka stage 8 (Tanaka 1976), which precedes major developmental processes, like dorsal closure and organogenesis. The results show that the expression levels of hypermethylated genes are significantly higher than those of hypomethylated genes when methylation occurs in all the studied regions, except in the 5’ UTR or in the exon of monoexonic genes (fig. 3B). It is worth noting, however, that the number of annotated 5’ UTRs and monoexonic genes are relatively low, which could explain the lack of significant differences between hypermethylated and hypomethylated gene expression.

### Hypermethylated genes show greater expression change higher than hypomethylated genes

Once again using the set of transcriptomes of Ylla et al. (2018), we examined the gene expression change between ED2 (when the peak expression of DNMT1 is already declining) and ED6 (4 days later) (fig. 1B). We determined the fold change (log_2_FC) of differentially expressed genes between these two stages, considering those having a | log_2_FC | ≥ 2 and false discovery rate of < 0.05. In this way, we identified 1,599 genes, 553 of which were hypermethylated and 1,046 of which were hypomethylated. As shown in figure 4A, both hypermethylated and hypomethylated genes increased or decreased their expression levels in similar proportions. Intriguingly, the change was less in hypermethylated genes, regardless of whether this change was incremental or decremental, and this was more significant when the methylation was in the introns (fig. 4B). This notion can be condensed by expressing the coefficient of variation (CV) of gene expression between ED2 and ED6, which is significantly lower in hypermethylated than in hypomethylated genes (fig. 4C). The data suggest that the expression change of hypermethylated genes has lower dispersion than that in hypomethylated genes. This led us to analyze the CV of the expression levels of each gene between biological replicates at the same stage.

**FIG. 4.**
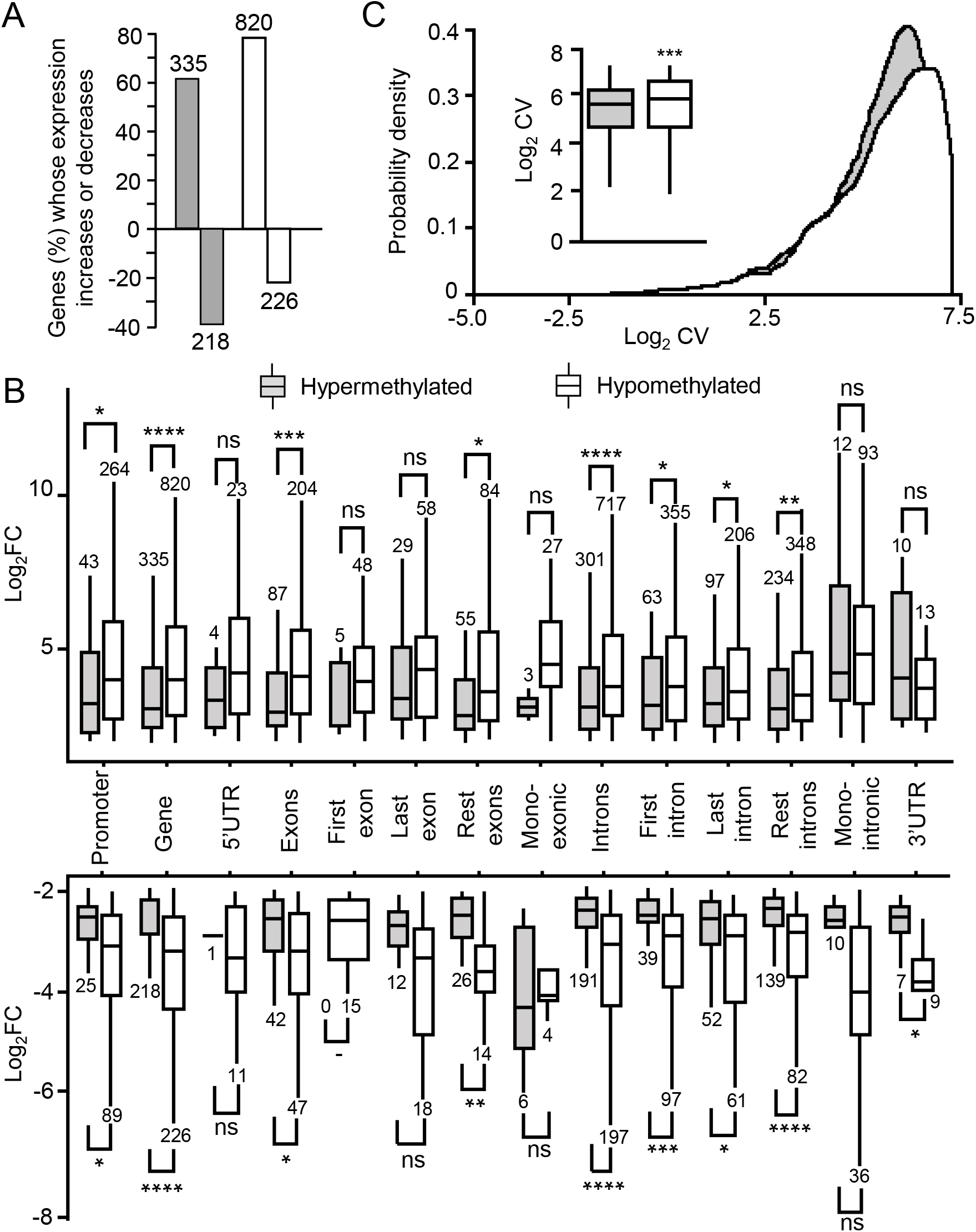
DNA methylation and gene expression dynamics in *Blattella germanica*. (A) Expression increase or decrease between ED2 and ED6 in hypermethylated and hypomethylated genes; a minimum of Log2FC > 2 with FDR < 0.05, was considered an increase or decrease. (B) Expression change (log_2_FC) between ED2 and ED6 of differentially upregulated or downregulated genes (Log_2_FC > 2 and FDR < 0.05); in all cases, genetic features were classified as hypermethylated or hypomethylated; data outliers have been omitted for clarity; the gene features considered were those in Table 1. (C) Density plot and boxplot (inset) of the coefficient of variation (CV) of the gene expression between ED2 and ED6, considering hypermethylated and hypomethylated genes; the inset describes the mean CV between ED2 and ED6 in hypermethylated and hypomethylated genes; data outliers have been omitted for clarity. In all cases, the grey bars indicate hypermethylated genes, and the white bars are hypomethylated genes; in B and C, asterisks indicate statistically significant differences using the Mann-Whitney U test, adjusting p-values by False Discovery Rate using the Benjamini-Hochberg method (* FDR < 0.05; ** FDR < 0.01; *** FDR < 0.001; **** FDR < 0.0001), non-significant differences (ns; FDR > 0.05), are also indicated.

### Hypermethylated genes have less expression variance than hypomethylated genes

Using two biological replicates, each comprising a pool of specimens from the transcriptome set of Ylla et al. (2018), we first compared the gene expression levels between replicates at the different stages. The results showed that there are no differences between replicates (supplementary fig. S2, Supplementary Material online). We then calculated the gene CV between the replicates in the same stage, comparing hypermethylated with hypomethylated genes at each stage. The results show that hypermethylated genes present less expression variability, measured as covariance, between biological replicates than hypomethylated genes do, a property that is more evident in earlier embryo stages (from ED0 to ED6) (fig. 5).

**FIG. 5.**
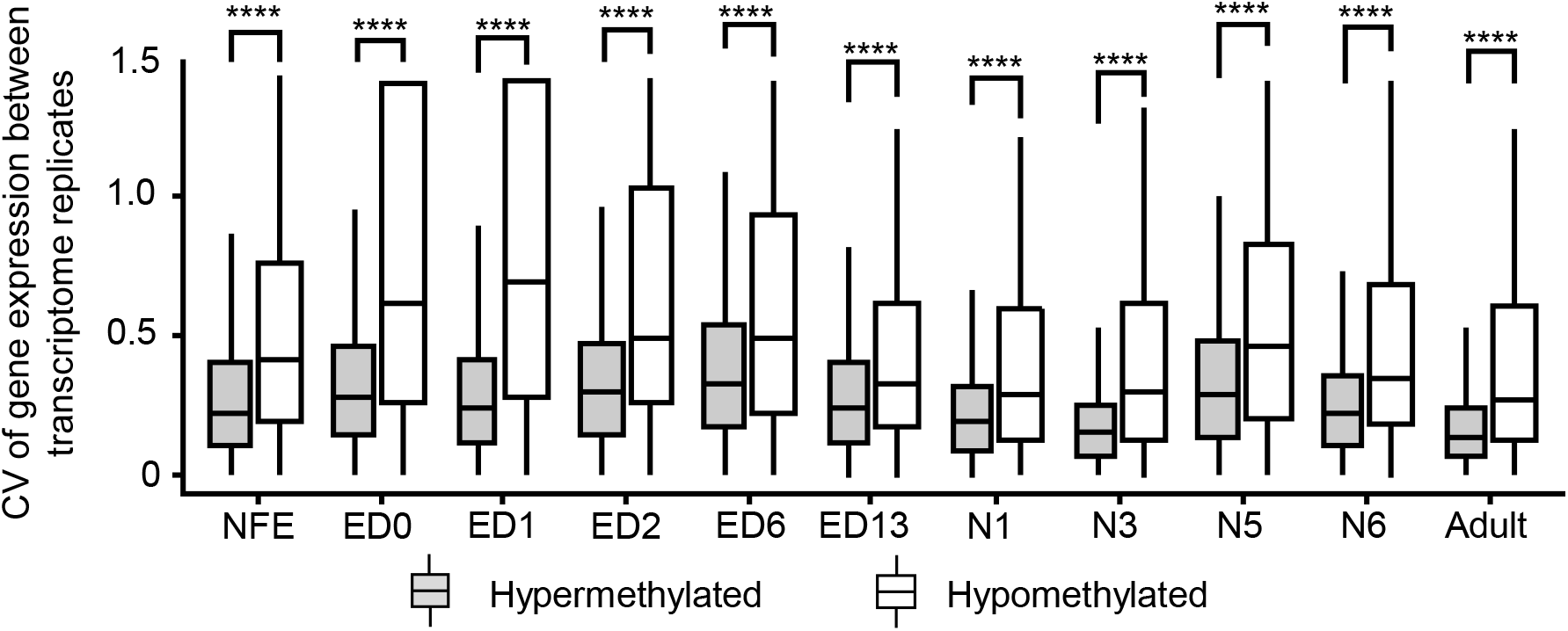
DNA methylation and expression variability in *Blattella germanica*. The coefficient of variation (CV) of gene expression for hypermethylated and hypomethylated genes between the two biological replicates generated from a pool of specimens dor each of the developmental transcriptomes studied. Data outliers have been omitted for clarity; the developmental stages studied were: NFE: non-fertilized egg; ED0 to ED13: embryo day 0 to embryo day 13; N1-N6: first to sixth nymphal stage; and the adult (Ylla et al. 2018). The four asterisks indicate statistically significant differences using the Mann-Whitney U test, adjusting p-values by False Discovery Rate using the Benjamini-Hochberg method (FDR < 0.0001).

## Discussion

The cockroach *B. germanica* has two *DNMT* genes, one coding for DNMT1 and one coding for DNMT3, which possess the functional motifs characteristic of these kinds of proteins, according to Lyko (2018). Quantitative determinations showed that *DNMT1* and *DNMT3* are expressed during the early embryo development (between 0% and 12% embryogenesis) of *B. germanica*. This suggests that both genes play roles in early embryogenesis, although *DNMT3* expression levels are about 100 times lower than those of *DNMT1*. To study these roles, we used maternal RNAi, which efficiently knocked down the *DNMT1*, but not the *DNMT3*, whose mRNA levels were not reduced despite the relatively high doses of dsRNA used, and the three independent experimental batches employed. Although *B. germanica* is highly sensitive to RNAi (Belles 2010), there are situations where this technique has been ineffective. For example, in the case of the lipophorin receptor, RNAi has proven highly efficient in the fatty body, where the gene is highly expressed, and much less efficient in the ovary, where it is expressed at low levels (Ciudad et al. 2007). In other cases, such as that of the yellow-g gene, the transience of its expression makes its depletion by RNAi impossible (Irles et al. 2009). We believe that the very low expression levels of *DNMT3* (about 1 copy of mRNA per 1000 copies of *Actin-5c* mRNA at most), are very difficult to significantly lower any further using RNAi.

In contrast, maternal RNAi of DNMT1was remarkably efficient. Indeed, the RNAi experiments and subsequent RRBS analyses showed that DNMT1 promotes DNA methylation in *B. germanica*, as observed in other insects, such as the milkweed bug *Oncopeltus fasciatus* (Bewick et al. 2019) and the beetle *T. castaneum* (Schulz et al. 2018), when implementing an equivalent approach. The RNAi experiments also revealed that DNMT1, and thus DNA methylation, is required to complete the germ band formation, in early embryogenesis, at 12% development. In the hymenopteran *Nasonia vitripennis* DNMT1-depleted embryos die at the onset of gastrulation (Zwier et al. 2012), at around 40% embryo development. In the beetle *T. castaneum*, although DNA methylation does not preferentially occur at CpG sites (Zemach et al. 2010; Feliciello et al. 2013; Song et al. 2017), DNMT1 is required in very early embryo development to progress beyond the first few cleavage cycles, in other words around 4% embryogenesis (Schulz et al. 2018). In the bug *O. fasciatus*, eggs laid by DNMT1-depleted females are inviable, although the stage at which development is interrupted has not been determined (Bewick et al. 2019). The fact that DNMT1 is required for embryo development in vertebrates, including mice (Li et al. 1992; Jackson-Grusby et al. 2001), frogs (Stancheva et al. 2001) and zebrafish (Rai et al. 2006), may suggest that their functions in embryogenesis are conserved from insects to vertebrates (Zwier et al. 2012). However, DNMT1 depletion in the insects *B. germanica*, *N. vitripennis*, and *T. castaneum* affects different embryo stages, and DNMT1 depletion in mice, frogs and zebrafish also elicits different phenotypes, resulting in the misexpression of genes that specify embryonic cell identity but with limited effects on early developmental mitosis (He et al. 2011). Thus, although DNA methylation is essential for embryogenesis in cockroaches up to mammals, the current evidence indicates that its specific action varies in different lineages, even within insects.

The matching expression patterns of *DNMT1* and *DNMT3* in *B. germanica* suggest that both act on the same early embryo stage, while the presence of a methyltransferase catalytic domain in DNMT1 and DNMT3 suggests that they both have the capacity to promote DNA methylation. However, when DNMT1 is depleted, a clear phenotype is observed in the embryo. This indicates that *DNMT3* expression, which is not affected by DNMT1 RNAi, does not compensate for the DNMT1 deficiency. These lines of evidence point to the possibility that both proteins are functionally redundant, and if so, *DNMT3*, which has very low expression levels, could be dispensable.

Most mammals have two DNMT3, DNMT3A and DNMT3B, which establish DNA methylation patterns. Even many rodent species have a third enzyme, DNMT3C that selectively methylate the promoters of young retrotransposon insertions in their germline (Molaro et al., 2020). In contrast, DNMT3 has been evolutionarily lost in a number of insect orders, including Odonata, Ephemeroptera, Orthoptera, Thysanoptera, Phthiraptera, Lepidoptera, Trichoptera, and Diptera (Bewick et al. 2017; Lewis et al. 2020b). Moreover, Bewick et al. (2017) showed that the presence of DNMT1 correlates positively with DNA methylation, whereas that is not seen for DNMT3. These authors suggest that either DNMT3 is unnecessary for DNA methylation or that DNMT1 compensates for DNMT3. Finally, studies in vitro have demonstrated that DNMT1 can also act as a de novo methylase (Fatemi et al. 2002). Taken together, the data suggest that *B. germanica* DNMT1 plays both the de novo and maintenance roles in DNA methylation, while DNMT3 has a minor role, and is possibly redundant with respect to DNMT1. It is worth noting that the DNMT3 sequence of *B. germanica*, especially the catalytic domain (supplementary fig. S3, Supplementary Material online), is remarkably conserved with respect to other proven functional DNMT3, such as that of the honeybee *A. mellifera* (Wang et al. 2006), suggesting that *B. germanica* DNMT3 is functional, and thus natural selection maintains the conserved sequence.

As in other species, the CG methylation levels in *B. germanica* present a bimodal distribution, being either very high or very low. In insects, this has also been reported in the locust *Schistocerca gregaria* (Falckenhayn et al. 2013) and the wasp *N. vitripennis* (Wang et al. 2013). Moreover, CG methylation in *B. germanica* tends to concentrate towards the 3’ region of the gene, in line with general DNA methylation trends in insects (Bewick et al. 2017; Lewis et al. 2020). In holometabolan species, DNA methylation appears to be biased towards the exons close to the 5’ region of the gene (Bonasio et al. 2012; Hunt et al. 2013; Wang et al. 2013; Glastad et al. 2016), while in hemimetabolans it presents higher levels towards the 3’ region of the gene coding part (Glastad et al. 2016; Bewick et al. 2019). In this sense, the DNA methylation pattern of hemimetabolans is similar to that of vertebrates, where the first intron and first exon are less methylated than the remaining regions in different tissues, species, and developmental stages (Anastasiadi et al. 2018). Furthermore, DNA methylation is biased towards exons rather than introns in some hemimetabolan insects, like the locust *S. gregaria* (Falckenhayn et al. 2013), the termite *Zootermosis nevadensis* (Glastad et al. 2016), and the bug *O. fasciatus* (Bewick et al. 2019), as is also the case in *B. germanica*. In general, our findings in embryos are similar to those observed by Bewick et al. (2019) in *B. germanica* adults, using whole-genome bisulfite sequencing data. These authors found similar general levels of CG methylation, with the highest being observed in intragenic regions rather than intergenic regions, and which tended to concentrate towards the 3’ UTR (Bewick et al. 2019; Lewis et al. 2020).

With respect to DNA methylation and gene functions, GO enrichment analyses revealed that hypermethylated genes are mainly involved in metabolic processes, and are more highly expressed than hypomethylated genes, which are instead related to signaling pathways. In other insects, like the ant *Camponotus floridanus* (Bonasio et al. 2012) and the wasp *N. vitripennis* (Wang et al. 2013), both holometabolan insects, hypermethylated genes are also enriched in housekeeping functions while hypomethylated genes are related to tissue-specific functions. Furthermore, it has recently been found that putatively methylated genes are under stronger purifying selection in both hemimetabolan and holometabolan insects (Ylla G, Nakamura T, Itoh T, Kajitani R, Toyoda A, Tomonari S, Bando T, Ishimaru Y, Watanabe T, Fuketa M, Matsuoka Y, Noji S, Mito T, Extavour CG, unpublished data, https://www.biorxiv.org/content/10.1101/2020.07.07.191841v1, last accessed August 25, 2020), highlighting the evolutionary importance of those genes undergoing DNA methylation.

A controversial aspect of DNA methylation is whether it can stimulate or repress gene expression. In vertebrates, a negative correlation between DNA methylation and gene expression has been reported, especially when methylation is located in the promoters, first intron, and first exon (Anastasiadi et al. 2018). In insects, a number of studies report a positive correlation between DNA methylation in intragenic regions and gene expression, such as in the termite *Z. nevadensis* (hemimetabolan) (Glastad et al. 2016), the ants *C. floridanus* and *Harpegnathos saltator* (Bonasio et al. 2012), and the wasp *N. vitripennis* (Wang et al. 2013) (holometabolans). However, in other insects like the locust *S. gregaria* (Falckenhayn et al. 2013), and the bug *O. fasciatus* (Bewick et al. 2019) (both hemimetabolans), no relationships have been found between DNA methylation and gene expression. Our observations indicate that hypermethylated genes are significantly more expressed than hypomethylated genes, especially in early embryogenesis (from ED0 to ED6), regardless of the methylation location in the gene.

The transcriptomic analysis of the hypermethylated and hypomethylated genes between ED2 and ED6 (i.e., after the *DNMT1* expression pulse) showed that the percentage of hypermethylated genes with increased expression was higher than the percentage of those with decreased expression (61% vs. 39%). Nevertheless, the hypomethylated genes behaved similarly (78% vs. 22%). Comparing the magnitude of change, in other words, how much the gene expression increased or decreased between ED2 and ED6, and the amount of expression variability (in terms of CV), revealed more exciting results. These results indicate that high DNA methylation levels were associated with high expression levels. At the same time, the amount of change between ED2 and ED6 was lower in hypermethylated genes, regardless of whether the change involved increasing or decreasing expression. This fits with the results of our GO enrichment analyses, as signaling factors are rarely expressed at high levels, but suffer higher expression variations, whereas high expression levels and low expression variation of housekeeping genes, and increased metabolism, is typical in early embryo development (Miyazawa and Aulehla 2018). Our results are reminiscent of those obtained for *N. vitripennis*, where methylated genes were shown to have higher median expression levels and lower expression variation across developmental stages than non-methylated genes (Wang et al. 2013).

Finally, comparing the gene expression between biological replicates at the different stages of *B. germanica* development, revealed that hypermethylated genes show lower expression variability than hypomethylated genes. This indicates that the expression of the methylated genes is tightly regulated, a feature that fits with the essential roles identified for these genes. The amount of intrinsic expression variability between individuals has been considered an inherent property of genes (Alemu et al. 2014; De Jong et al. 2019), representing a layer of gene regulation information that is just as important as changes in the mean expression levels (Wang and Zhang 2011). Expression variability has been shown to be low in genes involved in growth, general metabolism, and universal functions, whereas it is high in genes involved in environmental responses and non-housekeeping functions, in general, thus affecting gene network functioning by lowering noise (Alemu et al. 2014; De Jong et al. 2019).

Several features pertaining to the genomic, epigenomic, regulatory, polymorphic, functional, structural, and network characteristics of the gene, have been correlated with expression variability (Alemu et al. 2014). In the epigenomic context, a long-standing hypothesis posits that DNA methylation in gene regions reduces transcriptional noise, although the mechanisms involved are unclear (Bird 1995; Suzuki et al. 2007). However, there is scarce data supporting this hypothesis and it focuses on human tissues. Using nucleotide-resolution data on genomic DNA methylation and microarray data for human brain and blood tissues, Huh et al. (2013) showed that gene body methylation appears to lower expression variability. Further studies of human brain tissues, using Illumina sequencing, have indicated that genes with low expression variability are likely to have high gene methylation, whereas genes with low expression variability are likely to be non-methylated (Bashkeel et al. 2019). Our data on *B. germanica*, based on RRBS sequencing and transcriptomic data during embryo development, when DNA methylases are expressed and DNA methylation occurs, afford the first association between high DNA methylation and low expression variability in an insect.

The influence of DNA methylation on gene expression has aroused considerable interest, but the results from different models are disparate. For vertebrates and invertebrates, there is no consensus on whether DNA methylation stimulates or represses gene expression, nor on the influence of the methylated gene region. With regard to the phenotypic effects of DNA methylation, a representative number of studies on embryonic development in different vertebrate and invertebrate models could elucidate general trends. However, the effects reported are also disparate, since DNA methylation affects different embryogenetic processes in the various species studied, even within insects. Indeed, the only regularity that seems to operate in both vertebrates and invertebrates, involves the aforementioned association of high levels of DNA methylation with low expression noise. This generalization may even extend to plants, as suggested by recent studies of *Arabidopsis thaliana*, which indicate that genes with less expression variability are depleted in DNA methylation (Cortijo et al. 2019). It is true that the diversity of models studied in this sense is too small (a mammal, a cockroach, and a mustard plant) to draw definitive conclusions. However, the data provided by these models are very suggestive and could stimulate research on other species. Such research could eventually show that one of the few truly universal properties of DNA methylation is the reduction of expression noise.

## Materials and Methods

### Insects and dissections

Insects were obtained from a *B. germanica* colony fed with Panlab dog chow (Panlab S.L.U, Barcelona, Spain) and water *ad libitum*, and reared in the dark at 29±1°C and 60-70% relative humidity. Freshly ecdysed adult females were selected and used at appropriate ages. Mated females were used for all the experiments, and mating was assessed by checking the presence of sperm in the spermatheca. Prior to injection treatments, dissections, and tissue sampling, insects were anesthetized with carbon dioxide.

### Alignments and phylogenetic analysis of DNMT1 and DNMT3

Sequences used for the analyses were obtained by Blast from GenBank. Alignments were carried out with ClustalX (Larkin et al. 2007) and the phylogenetic tree reconstruction with 100 bootstraps of PhyML 3.1 (Guindon et al. 2010), based on the maximum-likelihood principle, a JTT matrix, a gamma model of heterogeneity rate, and using empirical base frequencies and estimating proportions. The sequences used for comparison with those of *B. germanica* were the following. For DNMT1, *Amyelois transitella* (XP_013186878.1), *Apis mellifera* (XP_026298868.1), *Athalia rosae* (XP_012254091.1), *Bombyx mori* (XP_012550860.1), *Danio rerio* (NP_571264.2), *Homo sapiens* (EAW84079.1), *Mus musculus* (EDL25141.1), *Nasonia vitripennis* (XP_008212391.1), *Nilaparvata lugens* (AHZ08393.1), *Tribolium castaneum* (XP_008193458.1) and *Zootermopsis nevadensis* (XP_021941799.1). For DNMT3 we used *A. mellifera* (XP_026302146.1), *D. rerio* (AAI62467.1), *H. sapiens* (3A1B_A), *M. musculus* (NP_001075164.1), *N. vitripennis* (XP_008204446.1) and *Z. nevadensis* (XP_021915977.1).

### RNA extraction and reverse transcription

Total RNA extraction was performed using RNeasy Plant minikit (QIAGEN, Hilden, Germany) in the case of young oothecae (from NFE to 4-day-old) and GenElute Mammalian Total RNA Miniprep kit (Sigma-Aldrich, Madrid, Spain) in the case of older oothecae (from 6- to 16-day-old). The volume extracted was lyophilized in a freeze-dryer FISHER-ALPHA 1–2 LDplus. Then, it was resuspended in 8 μL of miliQ H_2_O, treated with DNase I (Promega, Madison, WI, USA), and reverse transcribed with first Strand cDNA Synthesis Kit (Roche, Barcelona, Spain).

### Quantification of mRNA levels by quantitative real time PCR

Quantitative real-time PCR (qRT-PCR) reactions were carried out in triplicate in an iQ5 Real-Time PCR Detection System (Bio-Lab Laboratories, Madrid, Spain), using SYBR^®^Green (iTaq™ Universal SYBR^®^Green Supermix; Applied Biosystems, Foster City, CA, USA). A template-free control was included in all batches. Primers used to detect the transcripts are detailed in supplementary Table S1 (Supplementary Material online). The efficiency of each set of primers was validated by constructing a standard curve through three serial dilutions. mRNA levels were calculated relative to BgActin-5c mRNA (accession number AJ862721), using the Bio-Rad iQ5 Standard Edition Optical System Software (version 2.0). Results are given as copies of mRNA of interest per 1000 copies of BgActin-5c mRNA. To test the statistical significance between treated and control samples we used the Relative Expression Software Tool (REST), which evaluates the significance of the derived results by Pairwise Fixed Reallocation Randomization Test (Pfaffl 2002).

### RNA interference

Maternal RNAi assays have been described previously (Fernandez-Nicolas and Belles 2017). *DNMT1* and *DNMT3* sequences were amplified by PCR and then cloned into pSTBlue-1 vector. Primers used to prepare dsRNA are described in supplementary Table S1 (Supplementary Material online). A 307 bp sequence from *Autographa californica* nucleopolyhedrosis virus (Accession number K01149.1) was used as control dsRNA (dsMock). dsDNMT1 and dsDNMT3 were respectively injected at a dose of 3 μg in 1μl volume into the abdomen of 5-day-old adult females with a 5 μl Hamilton microsyringe. Control females were treated at the same age with the same dose and volume of dsMock.

### Microscopy

Twenty-day-old oothecae were detached from female abdomen, and artificially opened. Embryos were dechorionated and individualized. The embryos were directly observed under the stereo microscope Carl Zeiss–AXIO IMAGER.Z1 (Oberkochen, Germany). For 4’,6-di-amidino-2-phenylindole (DAPI) staining, 4-day-old oothecae were detached from female abdomen and incubated in PBT (Triton-X 0.1% in PBS 0.2M) in a water bath at 95°C. Then, each ootheca was artificially opened and embryos were dechorionated and individualized. Between 12 and 24 embryos per ootheca, chosen from the central part of it, were dissected for staining. The embryos were fixed in 4% paraformaldehyde in PBS 0.2M for 1 h, washed with PBT, and then incubated for 10 min in 1 mg/mL DAPI. They were mounted in Mowiol (Calbiochem, Madison, WL, USA) and observed with a fluorescence microscope Carl Zeiss–AXIO IMAGER.Z1.

### Reduced Represented Bisulfite Sequencing (RRBS) and methylation analyses

Four-day-old oothecae from dsDNMT1-treated females, and from controls (dsMock-treated) were detached from female abdomen and kept in a water bath at 95°C. Then, each ootheca was artificially opened and embryos were dechorionated and individualized. In the dsMock group, all embryos were collected. In the dsDNMT1 group, only embryos showing abnormal phenotypes (between 40 and 50%) were collected. Genomic DNA was extracted using GenElute™ Mammalian Genomic DNA Miniprep Kit (Merck), following manufacturer’s instructions. Then, samples were sent to the Genomics Unit of the Centre for Genomic Regulation (PRBB, Barcelona) where RRBS libraries were prepared using the Premium Reduced Representation Bisulfite Sequencing (RRBS) Kit (Diagenode), and sequenced ton an Illumina HiSeq 2000 platform in 50-bp single-end mode. We reserved a part of the sample to extract RNA, reverse transcribe, and check *DNMT1* transcript decrease by qRT-PCR. Sequences from RRBS libraries were first quality trimmed using Cutadapt (Marcel Martin 2011), and then aligned to the reference *B. germanica* genome (Accession code: PRJNA427252) using Bismark v.0.20.0. (Krueger and Andrews 2011). Once aligned, the bisulfite conversion ratio was calculated using spike-in controls provided by Premium Reduced Representation Bisulfite Sequencing (RRBS) Kit (Diagenode) (supplementary Table S2, Supplementary Material online). Then, using the methylKit package v1.4.1 for R we performed methylation calling; called bases with less than 10 reads or more than the 99th percentile of coverage were discarded. We checked that the four replicates have similar %mCG (supplementary fig. S4, Supplementary Material online). Using the mixtools R package, and assuming that DNA methylation follow a mixture of three different Gaussian distributions, we identified the Gaussian distribution of hypomethylated genes (mean methylation = 0.84 ± 0.99 %) and the Gaussian distribution of hypermethylated genes (mean methylation = 93.77 ± 4.45%). Then, we selected as hypermethylated regions those with methylation values between 80.43 - 100% and as hypomethylated regions those with methylation values between 0 – 3.82% (that is, using mean of each of the two peaks of the bimodal distribution of mCG ± 3 standard deviations to categorize the genes in hypermethylated and hypomethylated).

### Gene ontology (GO) analyses

Using gene annotations available at NCBI bioproject with accession number PRJNA427252, we assigned GO terms to *B. germanica* proteins using eggNOG mapper (Huerta-Cepas et al. 2017). Then, using topGO package (Alexa and Rahnenfuhrer 2010), we tested enriched GO terms in hypermethylated and hypomethylated genes. We used Fisher’s exact text with weighted algorithm to test statistical significance, which was stablished at p < 0.05.

### Combined analyses of genomic methylation and gene expression

For the transcriptomic analyses, we used RNA-seq libraries produced in our laboratory (Ylla et al. 2018), available at Gene Expression Omnibus with accession number GSE99785. To examine the changes in the gene expression between ED2 and ED6, we normalized raw counts using trimmed mean of M values (TMM), and performed a differential expression analysis between the embryo day 2 (ED2) and embryo day 6 (ED6) using the edgeR package (Robinson et al. 2009). Then, we selected the genes with an absolute log_2_FC ≥ 2 and FDR < 0.05 as differentially expressed genes. We separated the differentially expressed genes between upregulated and downregulated in ED6, and within each of these two groups we compared the expression levels of the hypomethylated vs hypermethylated genes with Mann–Whitney U test, adjusting p-values by false discovery rate (FDR) using Benjamini-Hochberg method. For the rest of the analyses we considered those genes having a | log_2_FC | ≥ 2 and FDR < 0.05. To test statistical differences between hypermethylated and hypomethylated genes FPKMs, | log_2_FC | and Coefficient of Variation (CV), we also used Mann–Whitney U test, adjusting p-values by FDR using Benjamini-Hochberg method. In the graphs, data outliers have been omitted for clarity. We defined as outliers those values below Q1-1.5*Inter Quartil Range (IQR) or higher than Q3+1.5*IQR, being IQR the distance between Q1 and Q3.

## Acknowledgments

This work was supported by the Spanish Ministry of Economy and Competitiveness (grants CGL2012-36251, CGL2015-64727-P and PID2019-104483GB-I00 to XB, including FEDER funds), the CSIC (grant 2019AEP029) and the Catalan Government (grants 2014 SGR 619 and 2017 SGR 1030).

## Author Contributions

X.B. designed the research; A.V.-A. X.B., J.C.M and G.Y. performed the research; GY and J.C.M performed the bioinformatics analyses; X.B., A.V.-A., G.Y. and J.C.M discussed and interpreted the results; A.V.-A. and X.B. wrote the paper.

## References

Alemu EY, Carl Jr JW, Corrada Bravo H, Hannenhalli S. 2014. Determinants of expression variability. Nucleic Acids Res. 42:3503–3514.

Alexa A, Rahnenfuhrer J. 2010. Bioconductor - topGO. R Packag. version 28.

Anastasiadi D, Esteve-Codina A, Piferrer F. 2018. Consistent inverse correlation between DNA methylation of the first intron and gene expression across tissues and species. Epigenet Chromatin 11(1):37.

Bashkeel N, Perkins TJ, Kærn M, Lee JM. 2019. Human gene expression variability and its dependence on methylation and aging. BMC Genomics 20:941.

Belles X. 2010. Beyond *Drosophila*: RNAi in vivo and functional genomics in insects. Annu. Rev. Entomol. 55:111–128.

Belles X. 2020. Insect metamorphosis. From natural history to regulation of development and evolution. London: Academic Press.

Bewick AJ, Sanchez Z, McKinney EC, Moore AJ, Moore PJ, Schmitz RJ. 2019. Dnmt1 is essential for egg production and embryo viability in the large milkweed bug, *Oncopeltus fasciatus*. Epigenetics & Chromatin 12:6.

Bewick AJ, Vogel KJ, Moore AJ, Schmitz RJ. 2017. Evolution of DNA methylation across insects. Mol. Biol. Evol. 34:654–665.

Bird AP. 1995. Gene number, noise reduction and biological complexity. Trends Genet. 11:94–100.

Bonasio R, Li Q, Lian J, Mutti NS, Jin L, Zhao H, Zhang P, Wen P, Xiang H, Ding Y, et al. 2012. Genome-wide and caste-specific DNA methylomes of the ants C*amponotus floridanus* and *Harpegnathos saltator*. Curr. Biol. 22:1755–1764.

Cardoso-Júnior CA, Fujimura PT, Santos-Júnior CD, Borges NA, Ueira-Vieira C, Hartfelder K, Goulart LR, Bonetti AM. 2017. Epigenetic modifications and their relation to caste and sex determination and adult division of labor in the stingless bee *Melipona scutellaris*. Genet. Mol. Biol. 40:61–68.

Ciudad L, Bellés X, Piulachs MD. 2007. Structural and RNAi characterization of the German cockroach lipophorin receptor, and the evolutionary relationships of lipoprotein receptors. BMC Mol. Biol. 8:53.

Cortijo S, Aydin Z, Ahnert S, Locke JC. 2019. Widespread inter-individual gene expression variability in *Arabidopsis thaliana*. Mol. Syst. Biol. 15:e8591.

Falckenhayn C, Boerjan B, Raddatz Gü, Frohme M, Schoofs L, Lyko F. 2013. Characterization of genome methylation patterns in the desert locust *Schistocerca gregaria*. J. Exp. Biol. 216:1423–1429.

Fatemi M, Hermann A, Gowher H, Jeltsch A. 2002. Dnmt3a and Dnmt1 functionally cooperate during *de novo* methylation of DNA. Eur. J. Biochem. 269:4981–4984.

Feliciello I, Parazajder J, Akrap I, Ugarković D. 2013. First evidence of DNA methylation in insect *Tribolium castaneum*: Environmental regulation of DNA methylation within heterochromatin. Epigenetics 8:534–541.

Fernandez-Nicolas A, Belles X. 2017. Juvenile hormone signaling in short germ-band hemimetabolan embryos. Development 144:4637–4644.

Glastad KM, Gokhale K, Liebig J, Goodisman MAD. 2016. The caste- and sex-specific DNA methylome of the termite *Zootermopsis nevadensis*. Sci. Rep. 6: 37110.

Glastad KM, Hunt BG, Goodisman MA. 2014. Evolutionary insights into DNA methylation in insects. Curr. Opin. Insect Sci. 1:25–30.

Goll MG, Kirpekar F, Maggert KA, Yoder JA, Hsieh CL, Zhang X, Golic KG, Jacobsen SE, Bestor TH. 2006. Methylation of tRNAAsp by the DNA methyltransferase homolog Dnmt2. Science. 311:395–398.

Guindon S, Dufayard JF, Lefort V, Anisimova M, Hordijk W, Gascuel O. 2010. New algorithms and methods to estimate maximum-likelihood phylogenies: Assessing the performance of PhyML 3.0. Syst. Biol. 59:307–321.

Harrison MC, Jongepier E, Robertson HM, Arning N, Bitard-Feildel T, Chao H, Childers CP, Dinh H, Doddapaneni H, Dugan S, et al. 2018. Hemimetabolous genomes reveal molecular basis of termite eusociality. Nat. Ecol. Evol. 2:557–566.

He XJ, Chen T, Zhu JK. 2011. Regulation and function of DNA methylation in plants and animals. Cell Res. 21:442–465.

Huerta-Cepas J, Forslund K, Coelho LP, Szklarczyk D, Jensen LJ, Von Mering C, Bork P. 2017. Fast genome-wide functional annotation through orthology assignment by eggNOG-mapper. Mol. Biol. Evol. 34:2115–2122.

Huh I, Zeng J, Park T, Yi SV. 2013. DNA methylation and transcriptional noise. Epigenetics & Chromatin 6:9.

Hunt BG, Glastad KM, Yi S V., Goodisman MAD. 2013. The function of intragenic DNA methylation: Insights from insect epigenomes. Integr. Comp. Biol. 53:319–328.

Irles P, Bellés X, Piulachs MD. 2009. Identifying genes related to choriogenesis in insect panoistic ovaries by Suppression Subtractive Hybridization. BMC Genomics 10:206.

Jackson-Grusby L, Beard C, Possemato R, Tudor M, Fambrough D, Csankovszki G, Dausman J, Lee P, Wilson C, Lander E, et al. 2001. Loss of genomic methylation causes p53-dependent apoptosis and epigenetic deregulation. Nat. Genet. 27:31–39.

Jones PA. 2012. Functions of DNA methylation: Islands, start sites, gene bodies and beyond. Nat. Rev. Genet. 13:484–492.

De Jong T V., Moshkin YM, Guryev V. 2019. Gene expression variability: The other dimension in transcriptome analysis. Physiol. Genomics 51:145–158.

Jurkowski TP, Meusburger M, Phalke S, Helm M, Nellen W, Reuter G, Jeltsch A. 2008. Human DNMT2 methylates tRNAAsp molecules using a DNA methyltransferase-like catalytic mechanism. RNA 14:1663–1670.

Krueger F, Andrews SR. 2011. Bismark: A flexible aligner and methylation caller for Bisulfite-Seq applications. Bioinformatics 27:1571–1572.

Larkin MA, Blackshields G, Brown NP, Chenna R, McGettigan PA, McWilliam H, Valentin F, Wallace IM, Wilm A, Lopez R, et al. Clustal W and Clustal X version 2.0. Bioinformatics 23:2947–2948.

Lewis SH, Ross L, Bain SA, Pahita E, Smith SA, Cordaux R, Miska EA, Lenhard B, Jiggins FM, Sarkies P. 2020. Widespread conservation and lineage-specific diversification of genome-wide DNA methylation patterns across arthropods. PLoS Genet. 16:e1008864.

Li B, Hou L, Zhu D, Xu X, An S, Wang X. 2018. Identification and caste-dependent expression patterns of DNA methylation associated genes in *Bombus terrestris*. Sci. Rep. 8:2332.

Li E, Bestor TH, Jaenisch R. 1992. Targeted mutation of the DNA methyltransferase gene results in embryonic lethality. Cell 69:915–926.

Lyko F. 2018. The DNA methyltransferase family: A versatile toolkit for epigenetic regulation. Nat. Rev. Genet. 19:81–92.

Martin M. 2011. Cutadapt removes adapter sequences from high-throughput sequencing reads. EMBnet 17:1–3.

Miyazawa H, Aulehla A. 2018. Revisiting the role of metabolism during development. Development 145: dev131110.

Molaro A, Malik HS, Bourc’his D. 2020. Dynamic evolution of de novo DNA methyltransferases in rodent and primate genomes. Mol. Biol. Evol. 37:1882–1892.

Pfaffl MW. 2002. Relative expression software tool (REST©) for group-wise comparison and statistical analysis of relative expression results in real-time PCR. Nucleic Acids Res. 30:e36.

Rai K, Nadauld LD, Chidester S, Manos EJ, James SR, Karpf AR, Cairns BR, Jones DA. 2006. Zebra fish Dnmt1 and Suv39h1 regulate organ-specific terminal differentiation during development. Mol. Cell. Biol. 26:7077–7085.

Robinson MD, McCarthy DJ, Smyth GK. 2009. edgeR: A Bioconductor package for differential expression analysis of digital gene expression data. Bioinformatics 26:139–140.

Sarda S, Zeng J, Hunt BG, Yi S V. 2012. The evolution of invertebrate gene body methylation. Mol. Biol. Evol. 29:1907–1916.

Schulz NKE, Wagner CI, Ebeling J, Raddatz G, Diddens-de Buhr MF, Lyko F, Kurtz J. 2018. Dnmt1 has an essential function despite the absence of CpG DNA methylation in the red flour beetle *Tribolium castaneum*. Sci. Rep. 8:16462.

Song X, Huang F, Liu J, Li C, Gao S, Wu W, Zhai M, Yu X, Xiong W, Xie J, et al. 2017. Genome-wide DNA methylomes from discrete developmental stages reveal the predominance of non-CpG methylation in *Tribolium castaneum*. DNA Res. 24:445–458.

Stancheva I, Hensey C, Meehan RR. 2001. Loss of the maintenance methyltransferase, xDnmt1, induces apoptosis in *Xenopus* embryos. EMBO J. 20:1963–1973.

Suzuki MM, Kerr ARW, De Sousa D, Bird A. 2007. CpG methylation is targeted to transcription units in an invertebrate genome. Genome Res. 17:625–631.

Tanaka A. 1976. Stages in the embryonic development of the German cockroach, *Blattella germanica* Linné (Blattaria, Blattellidae). Kontyû, Tokyo 44:1703–1714.

Wang X, Wheeler D, Avery A, Rago A, Choi JH, Colbourne JK, Clark AG, Werren JH. 2013. Function and evolution of DNA methylation in *Nasonia vitripennis*. PLoS Genet. 9:1003872.

Wang Y, Jorda M, Jones PL, Maleszka R, Ling X, Robertson HM, Mizzen CA, Peinado MA, Robinson GE. 2006. Functional CpG methylation system in a social insect. Science. 314:645–647.

Wang Z, Zhang J. 2011. Impact of gene expression noise on organismal fitness and the efficacy of natural selection. Proc. Natl. Acad. Sci. U. S. A. 108:E67–E76.

Ylla G, Piulachs M.D., Belles X. 2018. Comparative transcriptomics in two extreme neopterans reveals general trends in the evolution of modern insects. iScience 4:164–179.

Zemach A, McDaniel IE, Silva P, Zilberman D. 2010. Genome-wide evolutionary analysis of eukaryotic DNA methylation. Science 328:916–919.

Zwier M V, Verhulst EC, Zwahlen RD, Beukeboom LW, Van De Zande L. 2012. DNA methylation plays a crucial role during early *Nasonia* development. Insect Mol. Biol. 21:129–138.

