## Supplementary figures and Tables for "DNA methylation in cockroaches is essential in early embryo development and reduces gene expression noise"

Contents

**Figure S1.** Phylogenetic relationships of the DNMT1 and DNMT3 proteins of Blattella germanica with those of other insect species.

**Figure S2**. Violin plots displaying the FPKM distribution in log scale between the two transcriptome replicates of each studied developmental stage of *Blattella germanica*.

**Figure S3.** Alignment of the catalytic region of DNMT3 of *Blattella germanica* compared with that of *Apis mellifera*, *Nasonia vitripennis* and *Zootermopsis nevadensis.*

**Figure S4**. Percentage of CG methylation in the four RRBS libraries obtained in *Blattella germanica*.

**Table S1**. Primers used to measure the expression levels of DNMT1 and DNMT3 by qRT-PCR, and to prepare the corresponding dsRNAs for RNAi experiments.

**Table S2.** Statistics of the RRBS libraries, and bisulfite conversion ratio.

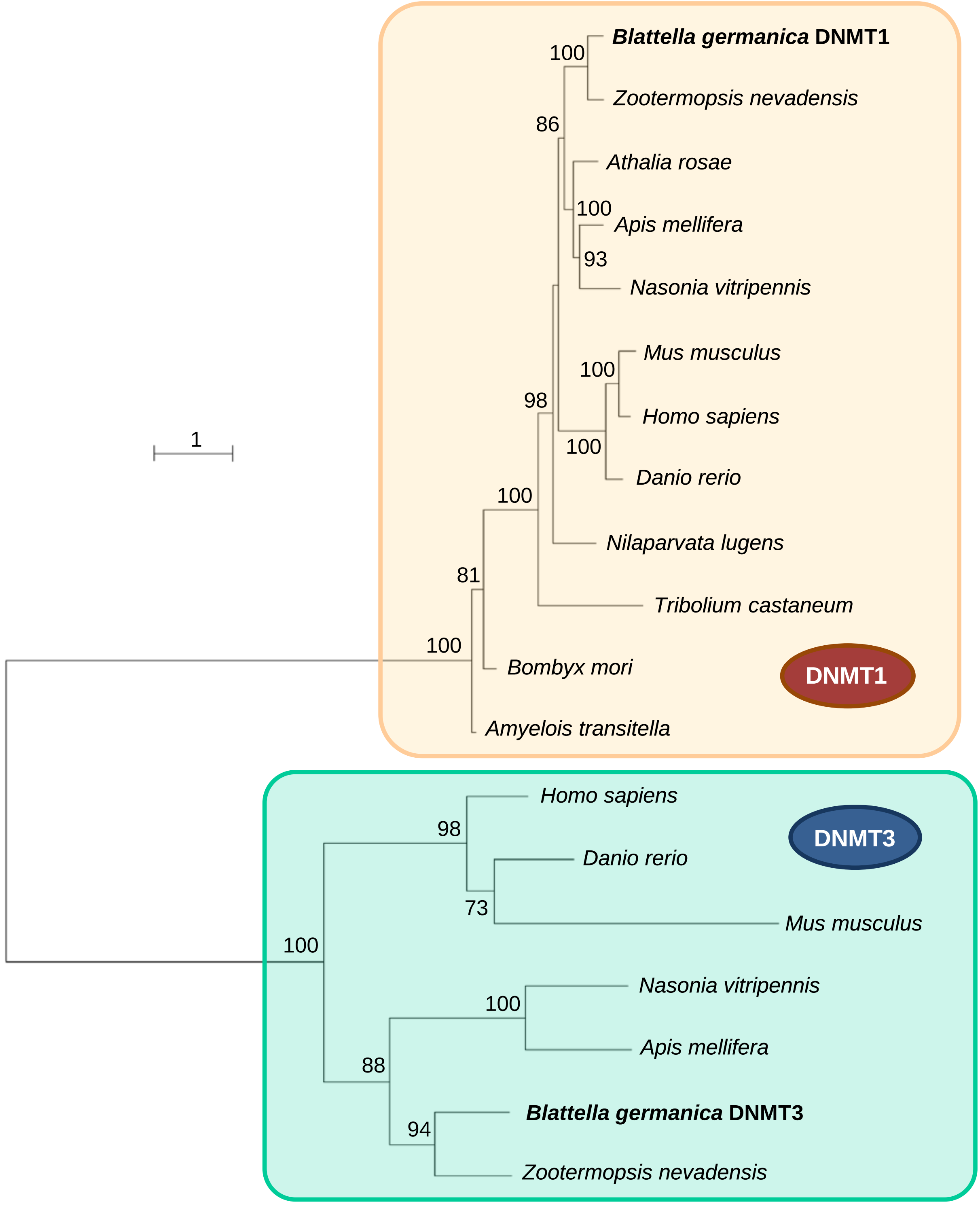


**Figure S1**. Phylogenetic relationships of the DNMT1 and DNMT3 proteins of *Blattella germanica* with those of other insect species. Sequences used were obtained by Blast from GenBank. Alignments were performed with ClustalX and phylogenetic reconstruction with PhyML 3.1 (http://www.atgc-montpellier.fr/download/papers/phyml_2010.pdf), based on the maximum-likelihood principle, a JTT matrix, a gamma model of heterogeneity rate, and using empirical base frequencies and estimating proportions. Data was bootstrapped for 100 replicates. The sequences used for comparison with those of *B. germanica* were the following. For DNMT1, *Amyelois transitella* (XP_013186878.1), *Apis mellifera* (XP_026298868.1), *Athalia rosae* (XP_012254091.1), *Bombyx mori* (XP_012550860.1), *Danio rerio* (NP_571264.2), *Homo sapiens* (EAW84079.1), *Mus musculus* (EDL25141.1), *Nasonia vitripennis* (XP_008212391.1), *Nilaparvata lugens* (AHZ08393.1), *Tribolium castaneum* (XP_008193458.1) and *Zootermopsis nevadensis* (XP_021941799.1). For DNMT3 we used *A. mellifera* (XP_026302146.1), *D. rerio* (AAI62467.1), *H. sapiens* (3A1B_A), *M. musculus* (NP_001075164.1), *N. vitripennis* (XP_008204446.1), *Z. nevadensis* (XP_021915977.1). Bootstrap values >50 are indicated in the corresponding nodes. Scale bar: number of substitutions per site.

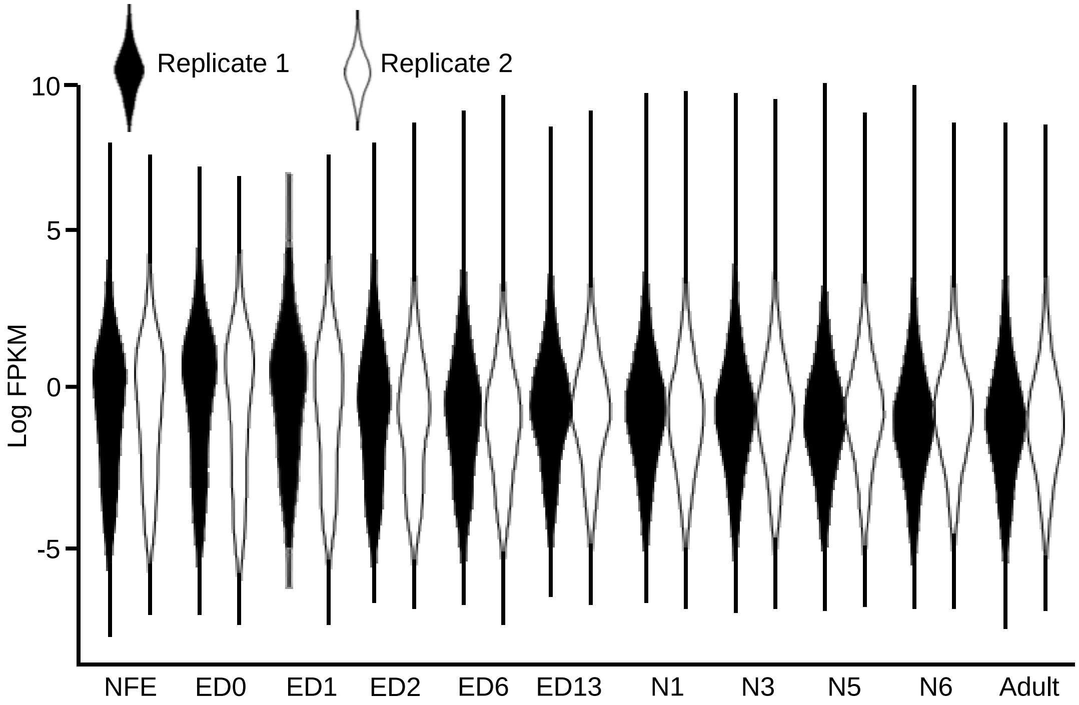


**Figure S2**. Violin plots displaying the FPKM distribution in log scale between the two transcriptome replicates of each studied developmental stage of *Blattella germanica*. The developmental stages are: NFE: non-fertilized egg, ED0 to ED13: embryo day 0 to embryo day 13, N1-N6: first to sixth nymphal stage, and the adult (Ylla et al., 2018).

```
Zootermopsis : -RVLSLFDGISTGMVVLKMMGIKVEKYYASEVDKDAINVSKVNHGDSIEHIGDVE
Blattella    : -TVLSLFDGIGTGLVVLKNLGIKVERYYASEVDVKAVEVCKFNHKEVIQ-IGDVT
Apis         : IRVLSLFDGLGTGLLVLLKLGFIVDAYYASEIDQDALMVTASHFGDRILQLGNVK
Nasonia      : IRVLSLFDGIGTGLVVLKHLNVNIECYYASEIDPDSMQVSFFNHGNEIIQLGDVR

Zootermopsis : LLSPEKLSRLGPIDLLIGGSPCTELSLVNPARKGLYDTEGSGYLFFDFYRVLMTL
Blattella    : KLPDNEISKL-QVDLLIGGSPCTELSLVNPARKGLFDTTATGFLFFEYYRILTIL
Apis         : DITCNTIKEIAPIDLLIGGSPCNDLSLANPARLGLHDPRGTGVLFFEYRRILKLV
Nasonia      : NIDEKKIKEIAPIDLLIGGSPCNELSLANPKRRGLDDPEGTGILFYDYVRIMKLV

Zootermopsis : QVL-HGKNIFWIFENTASMPYRLRNVITRFLGCDPVVIDASWLSAQRRARFFWGN
Blattella    : S---KKKTIFWLFENTAAMERTTRNTISRFFDRDPVVIDAVHFSAQRRARLFWGN
Apis         : RKLNNERHLFWLYENVASMPSEYRLEINKHLGQEPDVIDSADFSPQHRLRLYWHN
Nasonia      : KKHNKKRHLFWLFENVASMPKKFRNQISKNLGREPKFLDSADFSAQHRPRLYWGN

Zootermopsis : IPGLGRTELPEINMKLDECLMPGMERKAVVEKIRTVTTQSNSLLQGKNRIFPVKM
Blattella    : IPGLGYTNITDANFTLEQCLLPGFNRKAKVKKIRTVTSNPSSLLQGKTKNYPIDV
Apis         : FPIEPRLLSSQREQDVQDILTPHCQRYSLVKKIRTVTTKVNSLKQGKLALKPILM
Nasonia      : LPWGPYQVNN---VVLQDVLRKRCNRQALVKKIMTVTTRTNSLNQTKENLKPVLM

Zootermopsis : KEEMDAVWITELEVIFGLPLHYTDTGNLQLRERRQLLGRAWSVPVVKHILQPL
Blattella    : NGSPDRIWITELEMVFGFPIHYTDTGNINLQKRYKLLGKSWSVPAVQHILRP-
Apis         : KDESDSLWITELEEIFGFPRHYTDVKNLSATKRQRLIGKSWSVQTLTAIFESL
Nasonia      : DGKKDMLWVTELEKIFGFPMHYTDT-NLQKTRRLQLIGKAWSVQTLTAILR--
```

**Figure S3.** Alignment (Clustal X, Larkin et al., 2007) of the catalytic region of DNMT3 of *Blattella germanica* compared with that of *Apis mellifera* (XP_026302146.1), *Nasonia vitripennis* (XP_008204446.1) and *Zootermopsis nevadensis* (XP_021915977.1).

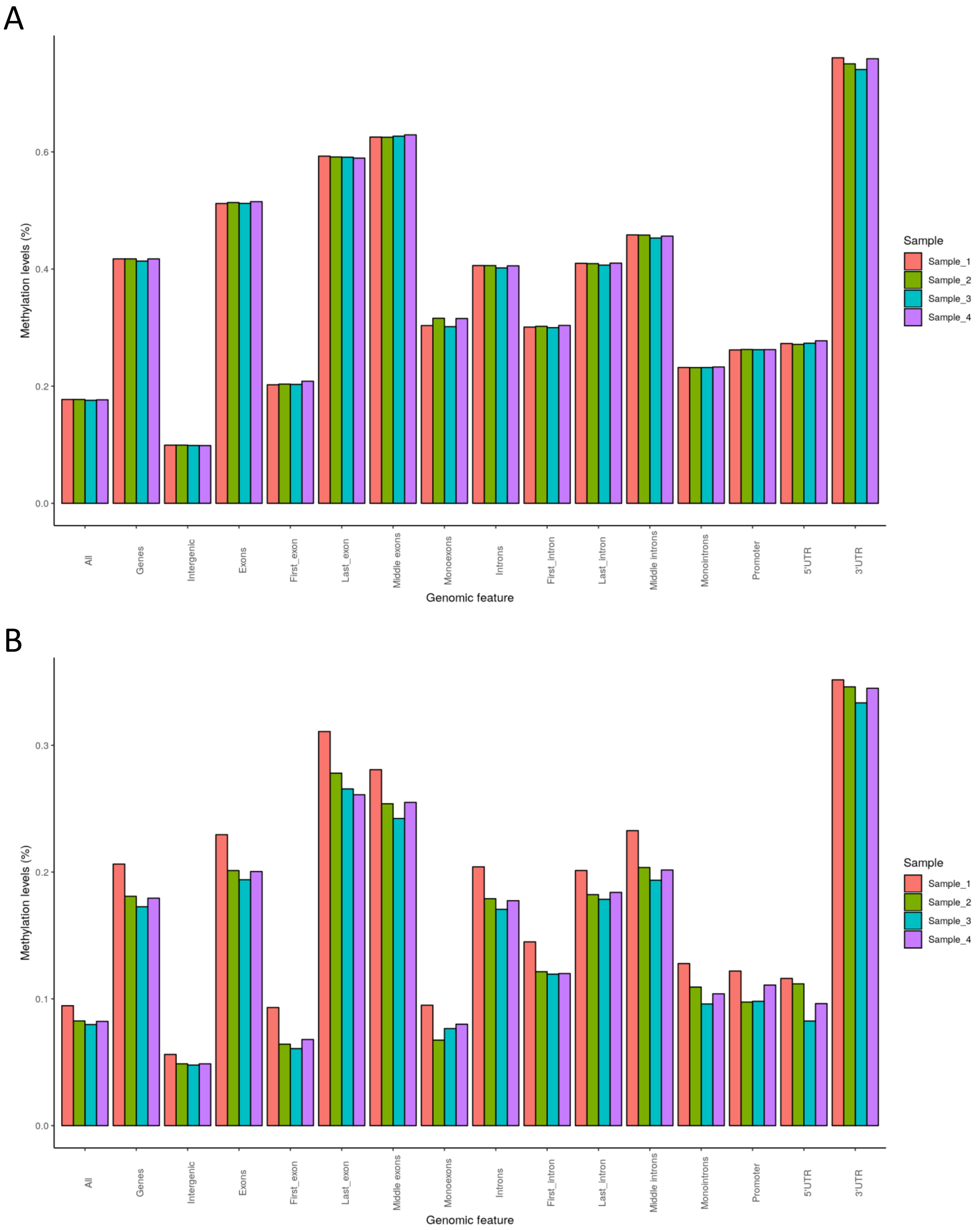


**Figure S4**. Percentage of CG methylation in the four RRBS libraries obtained in *Blattella germanica*. (A) RRBS of control insects. (B) RRBS of DNMT1-depleted insects. (%) levels in each biological replicate per every genomic feature studied. The genomic features examined are the same described in Table 1., The methylKit package v1.4.1 for R with methylation calling, was used for comparisons; called bases with less than 10 reads or more than the 99th percentile of coverage were discarded.

**Table S1**. Primers used to measure the expression levels of DNMT1 and DNMT3 by qRT-PCR, and to prepare the corresponding dsRNAs for RNAi experiments.

| Gene | Forward primer | Reverse primer | Accession number |
|---|---|---|---|
| Actin 5C | AGCTTCCTGATGGTCAGGTGA | TGTCGGCAATTCCAGGGTACATGGT | AJ862721.1 |
| DNMT1 (qRT-PCR) | ATGAAAAAGGCGGTTGTGAC | AAAAACGTCTTGCCGTCATC | MT881788 |
| DNMT1 (dsRNA) | GTCGGAGAAAGTGGCAAGAG | GTCACAACCGCCTTTTTCAT | MT881788 |
| DNMT3 (qRT-PCR) | TTGAAAACACGGCTGCTATG | TGTGCCGAAAAATGTACAGC | MT881790 |
| DNMT3 (dsRNA) | CGCCGAGCTAGGTTATTCTG | CAATCTTGGTGCTCTGAGTCC | MT881790 |

**Table S2.** Statistics of the RRBS libraries, and bisulfite conversion ratio. ED4C1-ED4C4 are the four libraries of DNMT1-depleted 4-day-old embryos, and ED4T1-ED4T4 the four control libraries.

| Library | ED4C1 | ED4C2 | ED4C3 | ED4C4 | ED4T1 | ED4T2 | ED4T3 | ED4T4 |
|---|---|---|---|---|---|---|---|---|
| Raw reads | 18,519,760 | 24,355,944 | 40,711,653 | 41,651,622 | 28,035,791 | 24,170,456 | 20,865,347 | 20,626,862 |
| Trimmed reads | 6,909,794 | 93,27,442 | 15,334,651 | 15,793,729 | 12,434,698 | 9,532,532 | 8,947,228 | 9,258,945 |
| Trimmed ratio (%) | 37.31 | 38.30 | 37.67 | 37.92 | 44.35 | 39.44 | 42.88 | 44.89 |
| Unique aligned reads | 9,061,483 | 12,677,679 | 20,339,272 | 21,053,385 | 10,820,311 | 11,012,600 | 8,698,668 | 7,621,916 |
| Unique alignment ratio (%) | 48.93 | 52.05 | 49.96 | 50.55 | 38.59 | 45.56 | 41.69 | 36.95 |
| Covered cytosines | 80,059,545 | 111,178,068 | 177,077,633 | 183,312,338 | 93,668,773 | 95,184,764 | 75,468,956 | 66,398,833 |
| Covered CpG | 19,942,568 | 27,626,895 | 44,141,130 | 45,914,155 | 23,151,959 | 23,545,622 | 18,702,854 | 16,430,827 |
| CpGs > 10 reads | 563,909 | 722,180 | 818,622 | 816,939 | 531,916 | 530,875 | 451,577 | 412,469 |
| Fold coverage | 28.94 | 32.7 | 47.9 | 50.34 | 38.35 | 38.86 | 36.45 | 34.63 |
| Bisulfite conversion ratio (%) | 98.4 | 98.4 | 98.7 | 98.7 | 98.4 | 98.4 | 98.2 | 97.6 |
